# HSV-1 ICP22 is a selective viral repressor of cellular RNA polymerase II-mediated transcription elongation

**DOI:** 10.1101/2021.06.08.447513

**Authors:** Nur F. Isa, Olivier Bensaude, Nadiah C. Aziz, Shona Murphy

## Abstract

The Herpes Simplex Virus (HSV-1) immediate early protein ICP22 interacts with cellular proteins to inhibit host cell gene expression and promote viral gene expression. ICP22 inhibits phosphorylation of Ser2 of the RNA polymerase II (pol II) carboxyl-terminal domain (CTD) and productive elongation of pol II. Here we show that ICP22 affects elongation of pol II through both the early-elongation checkpoint and the poly(A)-associated elongation checkpoint on a protein-coding gene model. Coimmunoprecipitation assays using tagged ICP22 expressed in human cells and pulldown assays with recombinant ICP22 *in vitro* coupled with mass spectrometry identify transcription elongation factors, including P-TEFb, additional CTD kinases and the FACT complex as interacting cellular factors. Using a photoreactive amino acid incorporated into ICP22, we found that L191, Y230 and C225 crosslink to both subunits of the FACT complex in cells. Our findings indicate that ICP22 physically interacts with critical elongation regulators to inhibit transcription elongation of cellular genes, which may be vital for HSV-1 pathogenesis. We also show that the HSV viral activator, VP16 has a region of structural similarity to the ICP22 region that interacts with elongation factors, suggesting a model where VP16 competes with ICP22 to deliver elongation factors to viral genes.

## Introduction

Suppression of host gene expression during HSV-1 productive infection may not only dampen host immune responses but also increase the cellular resources available for viral gene expression, and requires interaction between viral and host factors. Expression of HSV-1 ICP22, one of five immediate early genes, is initiated by the viral tegument transcription activation protein VP16 in the absence of *de novo* viral protein synthesis (1, 2). Deletion of ICP22 from HSV-1 causes a significant decrease in viral replication and the establishment of latency (3, 4). While ICP22 helps to promote expression of HSV-1 late viral genes in productive infection, this protein has the opposite effect on host gene and early viral gene expression (5-8). Based on the amino acid identity and sequence conservation of ICP22 and its homologues in other herpesviruses, Schwyzer et al. (1994) proposed that ICP22 has four distinctive domains (9). The first 160 residues of the N-terminal region and the C-terminal residues 398-420 of ICP22 are unique to HSV-1 and HSV-2 (9, 10). The second domain of ICP22 from residues 161-292 is described as the core conserved region (CCR) and has high similarity among the human and animal alphaherpesviruses (9, 11). The third region comprises residues 293-386 and is an acidic domain containing serine and threonine residues. At the C-terminal end is a basic domain containing proline residues (9). The N- and C-terminal and acidic domains of ICP22 and its homologues vary in length, and these domains therefore give rise to unique ICP22 proteins within the herpesviruses.

Several studies indicate that ICP22 physically interacts or colocalises with host nuclear proteins (12-15). A short sequence within the ICP22^CCR^ (residues 193-256) is sufficient for interaction with and inhibition of P-TEFb, a cyclin-dependent kinase complex that phosphorylates the RNA polymerase II (pol II) carboxyl-terminal domain (CTD) (11). The pol II CTD consists of heptapeptide repeats of Tyr1-Ser2-Pro3-Thr4-Ser5-Pro6-Ser7, which are highly conserved in eukaryotes (16, 17). The CDK9 subunit of P-TEFb phosphorylates NELF-E subunit of negative elongation factors NELF, SPT5 subunit of DRB sensitivity-inducing factor DSIF, and Ser2 of the pol II CTD, which all together enables pol II to overcome a promoter-proximal early-elongation checkpoint (EEC) to promote the transition of pol II into productive elongation. P-TEFb regulates recruitment of the PAF1 elongation complex (subunits PAF1, LEO1, CTR9, CDC73 and WDR61), which is required for pol II pause release and transcription elongation through chromatin (18, 19). The P-TEFb-associated elongation factors FACT (subunits SSRP1 and SPT16) have also been shown to facilitate pol II elongation through nucleosomes and acts cooperatively with P-TEFb to alleviate inhibition by DSIF and NELF (20, 21). In addition to elongation complexes, CTD kinases such as CDK12, which is also a Ser2 pol II CTD kinase that shares structural similarity to P-TEFb (22, 23), add another layer of complexity to the transcriptional regulatory network. The activity of P-TEFb is highly regulated; it is sequestered in a catalytically-inactive 7SK snRNP, comprising 7SK snRNA, HEXIM, MEPCE and LARP7 and phosphorylation of Thr186 in the activation segment is required for CDK9 kinase activity (24-26) . We have shown that P-TEFb activity is also required for elongation through the poly(A)-associated checkpoint at the 3’ end of human protein-coding genes (27). Pol II is unable to go through these checkpoints and prematurely terminates when cells are treated with CDK9 inhibitors, like DRB (5,6-dichlorobenzimidazone-1-β-D-ribofuranoside).

In order to gain a better understanding of the mechanism of action of ICP22, we have explored the interaction of ICP22 with host cell proteins using *in vivo* and *in vitro* approaches. Here we confirm that the ICP22^CCR^ has the same effect on elongation of transcription by pol II through checkpoints at the beginning and end of genes as inhibition of P-TEFb by small molecules like DRB. We also show through a combination of pulldown-mass spectrometry approaches, that ICP22 interacts with several crucial pol II transcription elongation factors. Using amber suppression and UV crosslinking to ectopically-expressed ICP22 in cells, we show that ICP22 interacts directly with the FACT elongation complex.

Our findings emphasize that ICP22 acts as a selective viral repressor at the level of transcription elongation of human genes.

## Results

### ICP22 inhibits pol II elongation at the beginning and end of host cell genes

We have investigated the ability of the CCR of ICP22 to induce loss of pol II CTD Ser2 phosphorylation, a mark indicative of transcription elongation, by monitoring the effect of ectopic expression of Myc-ICP22^CCR^ on Ser2P in a time-course experiment (Fig. 1A). Myc-ICP22^CCR^ is localised in the nucleoplasm and detectable as early as 6 hours post-transfection (hpt) by western blot. The level of Ser2P decreases at 8 hpt of Myc-ICP22^CCR^ with little effect on the total pol II level, indicating specific loss of CTD phosphorylation. To understand how ICP22 affects the control of transcription elongation on host genes, we carried out pol II and Ser2P pol II CTD chromatin immunoprecipitation (ChIP)-qPCR on a model cellular gene, *KPNB1* (Fig. 1B) at 7 hpt of Myc-ICP22^CCR^ as this time point should provide a snapshot of the repressive action of ICP22 on actively-transcribing pol II. The level of pol II (Fig. 1C; Pol II) at the beginning of *KPNB1* is significantly elevated at the TSS (TSS+0.9) compared to that of control and mock, suggesting increased pol II pausing near the EEC. In addition, the level of pol II is significantly decreased downstream of the TSS (TSS+5.3), suggesting that the newly-initiated pol II is unable to make the transition to productive elongation. The level of pol II in untreated, mock and transfected cells in the gene body (TSS+20.2, pA-4.8 and pA-2.9) remains unaffected, suggesting that pol II that has passed the early elongation checkpoint before or during early accumulation of Myc-ICP22^CCR^ is able to continue elongation of this 36 kb-long gene. The level of pol II is significantly reduced downstream of the poly(A) site (pA-0.4, pA+1.4, pA+2.6 and pA+4.1) suggesting that Myc-ICP22^CCR^ affects the level of transcribing pol II at the end of *KPNB1*. ChIP analysis of Ser2P (Fig. 1C; Ser2P) indicates that the level of this mark decreases downstream of the TSS (TSS+5.3 and TSS+20.2) and of the poly(A) site (pA-0.4, pA+1.4, pA+2.6 and pA+4.1) of *KPNB1*. When normalized to the level of pol II (Fig. 1C; Ser2P/pol II), Ser2P levels are lower at the beginning and end of the gene but unaffected in the *KPNB1* gene body. Although the global level of Ser2P, as measured by western blot, remains unchanged at 7 hpt of Myc-ICP22^CCR^ (Fig. 1A; Ser2P/pol II intensity), the level of Ser2P is clearly affected on the gene. The level of pol II is unaffected at 6 hpt of Myc-ICP22^CCR^ (Fig. 1D; 6 h), while significant pausing of pol II at the TSS and loss of pol II in the gene body is observed at 8 hpt of Myc-ICP22^CCR^ (Fig. 1D; 8 h). Although ICP22 affects CTD phosphorylation and pol II elongation, the level of the KPNB1 protein is not affected (Supplementary Fig. S1), most likely due to the short ICP22 expression times. Accordingly, we conclude that ICP22 causes a failure of pol II to negotiate the early elongation checkpoint at the beginning of *KPNB1* gene and to induce premature termination of elongating pol II at a checkpoint close to the poly(A) site at the end of *KPNB1* gene, in the same way as the CDK9 inhibitors KM05382 and DRB (27). These results are in line with previous findings that ICP22 inhibits P-TEFb activity (11, 28). ICP22 may also inhibit other Ser2 pol II CTD kinases, including CDK12 which is also a critical activator of pol II transcription elongation (22, 29).

**Fig. 1:**
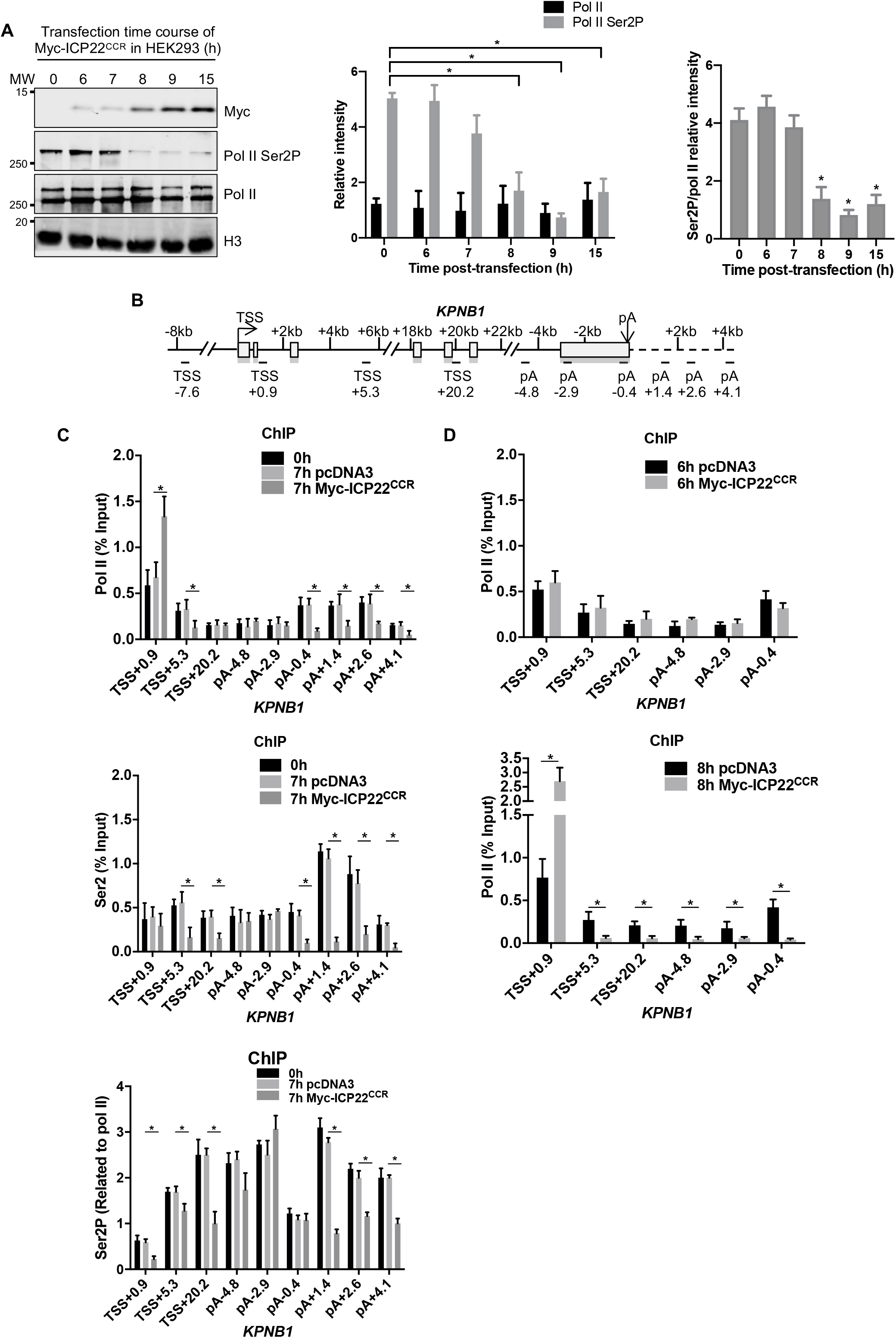
ICP22 inhibits pol II elongation at the beginning and end of host cell gene. A. HEK293 cells were transfected with Myc-ICP22^CCR^ and harvested at the indicated time. Nuclear lysates were analysed by western blot with Myc tag, Ser2P pol II CTD, pol II and H3 antibodies. H3 serves as the loading control. The relative intensity of protein signal of pol II, Ser2P and ratio of Ser2P/pol II were calculated from three independent experiments. Error bars indicate SEM and asterisks indicate the statistically significant change compared with 0 h (p<0.05). B. Schematic of *KPNB1* indicated in kilobase pairs (kb), with the transcription start site (TSS) and poly(A) site (pA) marked by arrows, exons by boxes and introns by lines (27, 29). The region of *KPNB1* amplified in quantitative PCR (qPCR) after chromatin immunoprecipitation (ChIP) is indicated underneath the gene. The ChIP-qPCR signal of each region is normalized to TSS-7.6, an upstream region with no detectable pol II to facilitate comparison of the level of factor associated with the different regions of the *KPNB1* gene. C. qPCR analysis of pol II and Ser2P ChIP after 7 h of transfection of pcDNA3 or Myc-ICP22^CCR^ in HEK293. Error bars indicate the SEM of n = 3 independent experiments and asterisks indicate the statistical significance compared with 7 h pcDNA3 (p<0.05). D. qPCR analysis of pol II and Ser2P ChIP after 6 h (top) and 8 h (bottom) of transfection of pcDNA3 or Myc-ICP22^CCR^ in HEK293. Error bars indicate the SEM of n = 3 independent experiments and asterisks indicate the statistical significance compared with 6 h or 8 h pcDNA3, respectively (p<0.05).

### ICP22 interacts with cellular transcription elongation factors

Having confirmed that ICP22^CCR^ inhibits pol II transcription elongation, we next addressed what transcription elongation factors interact with ICP22 using coimmunoprecipitation of full-length ICP22 and ICP22^CCR^ ectopically expressed in HEK293 cells. The proteins that copurify with ICP22 were analysed by western blot (Fig. 2A). CDK9/Cyclin T1, CDK12 and its cyclin, Cyclin K, PAF1 elongation complex components PAF1, LEO1 and CDC73 and the P-TEFb substrate SPT5 copurify with both full length ICP22 and ICP22^CCR^. In agreement with previous work, CDK7 does not copurify with ICP22 (15, 30). These results suggest that ICP22 interacts with CDK12 in addition to P-TEFb and with large elongation complexes that include the PAF1 complex.

**Fig. 2:**
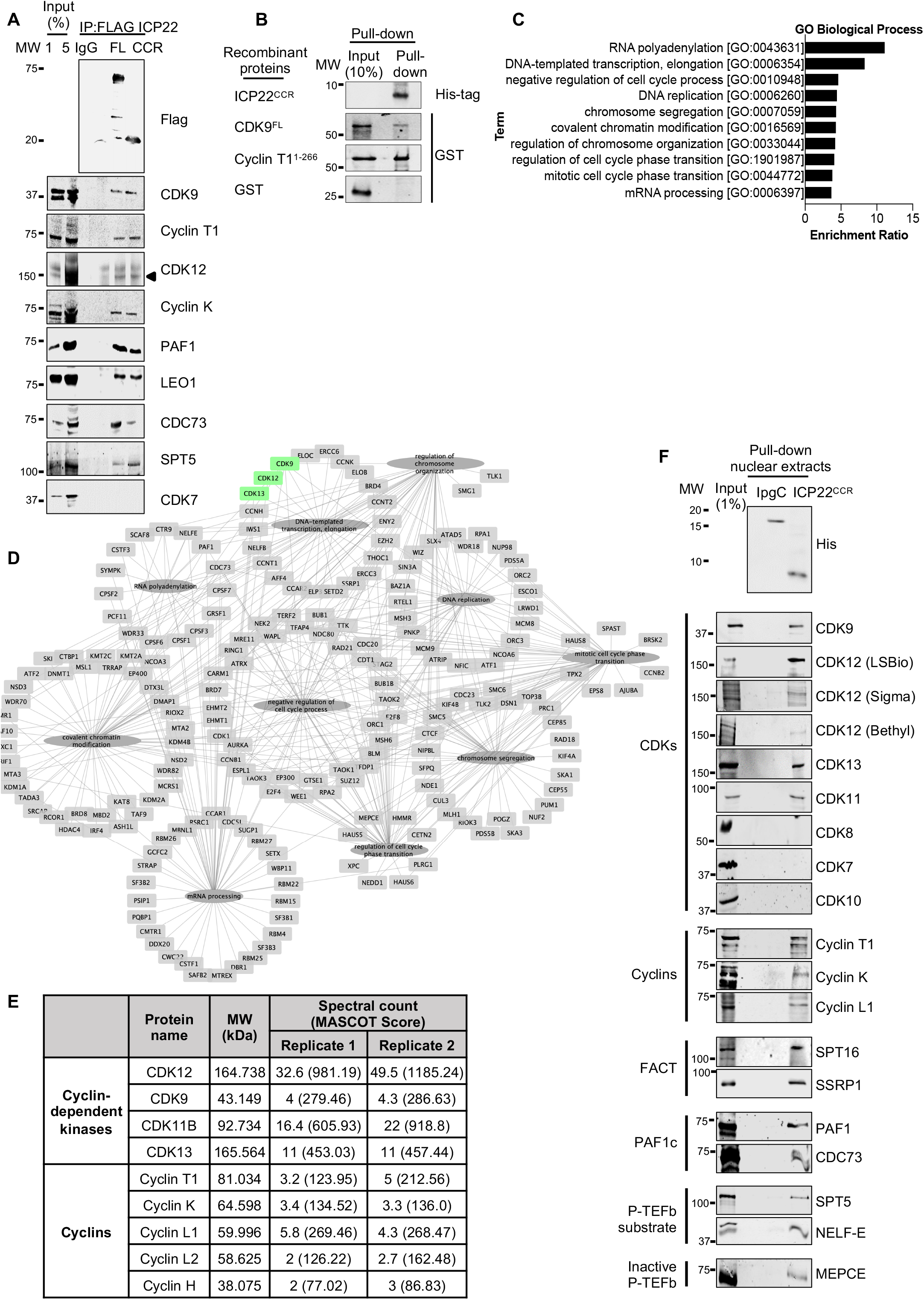
ICP22 interacts with cellular transcription elongation factors. A. HEK293 cells were transiently expressed with Flag-ICP22, either full length (FL) or the CCR. Cells were lysed, nuclear lysates were carefully separated for IP assays using control IgG or anti-Flag beads. Copurified proteins were determined by western blot using the antibodies noted at the right. B. His-ICP22^CCR^ was bacterially expressed and immobilized on Ni-NTA beads. GST-Cyclin T1^1-266^ and GST-CDK9^FL^ were expressed separately in *E*.*coli*. Immobilized GST-Cyclin T1^1-266^ and GST-CDK9^FL^ were eluted with reduced glutathione, and then incubated individually with His-ICP22^CCR^. Both input and proteins retained on the beads after washes were subjected to western blot with GST and His-tag antibodies as noted. C. Bar chart of the top ten gene ontology (GO) of biological processes set with a false discovery rate (FDR) ≤ 0.05 for proteins identified by mass spectrometry from His-ICP22^CCR^ pulldowns with HeLa nuclear extracts. The enrichment ratio indicates the number of observed genes divided by the number of expected genes in each GO category. D. Graphical representation of ICP22^CCR^ interacting proteins (represented by the official gene name) as categorised by GO terms, constructed using Cytoscape (version 3.5.1). Centrally positioned circular nodes indicate the description of the GO term, the surrounding square nodes denote the interacting proteins identified by mass spectrometry and the edges show biological processes associated with each protein. CDK9, CDK12, and CDK13 identified from GO term analysis are highlighted in green. E. Mass spectrometric identification of CDKs and cyclins copurified with ICP22^CCR^. F. His-ICP22^CCR^ pulldowns with HeLa nuclear extracts were determined by western blot using the antibodies noted on the right. The class of complex that the protein belongs to is noted on the left. His-IpgC which does not bind to P-TEFb was used as comparison. Antibodies against CDK12 from different manufacturers were used as indicated.

As P-TEFb comprises a heterodimeric cyclin–CDK complex, we tested the different subunits for interaction with ICP22. Recombinant His-tagged ICP22^CCR^, GST-CDK9^FL^, GST-Cyclin T1^1-266^ and GST alone, which serves as a control, were expressed and purified from bacteria and used for pulldown assays by incubating the ICP22^CCR^ with each P-TEFb subunit (Fig. 2B). His-ICP22^CCR^ weakly interacts with GST-CDK9 but interacts well with GST-Cyclin T1^1-266^. ICP22 is therefore likely to interact directly with both subunits of P-TEFb and residues 193-256 are sufficient to interact with P-TEFb subunits. In addition, P-TEFb complex formation and mammalian cell-specific post-translation modifications of the P-TEFb subunits are not required for interaction.

To identify any additional interacting proteins, we carried out a pulldown assay using bacterially-expressed recombinant His-ICP22^CCR^ and HeLa cell nuclear extracts. To control for the specificity of binding, *Shigella flexneri* chaperone protein IpgC, a small protein with comparable molecular mass to ICP22^CCR^, which is not known to directly interact with human proteins, with an N-terminal His-tag, was used as negative control (31). Importantly, *S. flexneri* IpgC does not interact with CDK9 nor Cyclin T1 (Supplementary Fig. S2). ICP22^CCR^ and IpgC interaction partners were analysed by mass spectrometry. We compiled a list of ICP22-interacting proteins with at least two unique peptides per protein that were detected in two independent proteomic experiments for further analysis (extended list of proteins in Supplementary Table 1). Using WebGestalt for Gene Ontology (GO) analysis, the top ten GO terms that are significantly associated with our ICP22 interactome are RNA polyadenylation and transcription elongation, consistent with the expectation that ICP22 affects host transcription elongation. (Fig. 2C and Supplementary Table 2). The ICP22 interactome includes several transcriptional kinases and their cyclin partners; CDK9/CyclinT1, CDK12/Cyclin K, CDK13/Cyclin K, CDK11B/Cyclin L1 or Cyclin L2 (Fig. 2D-E). Cyclin H is identified by mass spectrometry but not its known partners CDK7 and CDK20. P-TEFb-associated interactors such as PAF1, NELF-E, FACT complex subunits SPT16 and SSRP1, and SPT5 are present in both proteomic replicates. Western blot analysis confirms interaction of ICP22^CCR^ with CDK9, CDK12, CDK13, and CDK11 but not with CDK8, CDK7 and CDK10, validating identification of the ICP22-interacting proteins by mass spectrometry (Fig. 2F). Additionally, we validated the association of ICP22^CCR^ with cyclins (Cyclin T1, Cyclin K and Cyclin L1), the FACT complex (SPT16 and SSRP1), the PAF1 complex (PAF1 and CDC73), P-TEFb substrates SPT5 and NELF-E and a component of the P-TEFb inactive complex (MEPCE). We conclude that the coimmunoprecipitation of endogenous protein complexes and the combined approaches of pulldown-mass spectrometry performed in this study have clearly demonstrated the interaction of ICP22 with the host transcription elongation apparatus. On the basis of these results, we propose that ICP22 acts not only through interaction with and inhibition of CTD kinases but also through association with other elongation factors including the chromatin-specific transcription elongation FACT complex.

### UV crosslinking reveals that ICP22 interacts directly with the FACT complex

We next employed an unbiased approach where a UV-photo-crosslinkable amino acid is genetically incorporated into Flag-ICP22 for expression in HEK293 cells, followed by live-cell UV irradiation to induce crosslinking, coimmunoprecipitation in stringent buffer and mass spectrometry to detect cellular proteins that crosslink to ICP22. To identify cellular factors interacting with the ICP22^CCR^, several residues between 193-256 and three residues just outside this region (W187, L191 and W262) were replaced by an amber stop codon (TAG) in N-terminal Flag-tagged ICP22 (Fig. 3A). HEK293 were cotransfected with the mutant ICP22, suppressor tRNA (tRNA^Sup^) and aminoacyl tRNA synthetase specific for both the tRNA^Sup^ and the unnatural photoreactive amino acid p-benzoylphenylalanine (Bpa). Expression of ICP22^Bpa^ was optimized to produce photo-crosslinkable ICP22 analysed by anti-Flag western blot. Only truncated ICP22 is produced in the absence of Bpa, whereas full-length ICP22 that migrates with a molecular weight 60-75 kDa is successfully expressed for a range of mutants when Bpa is added to the culture medium (Fig. 3B).

**Fig. 3:**
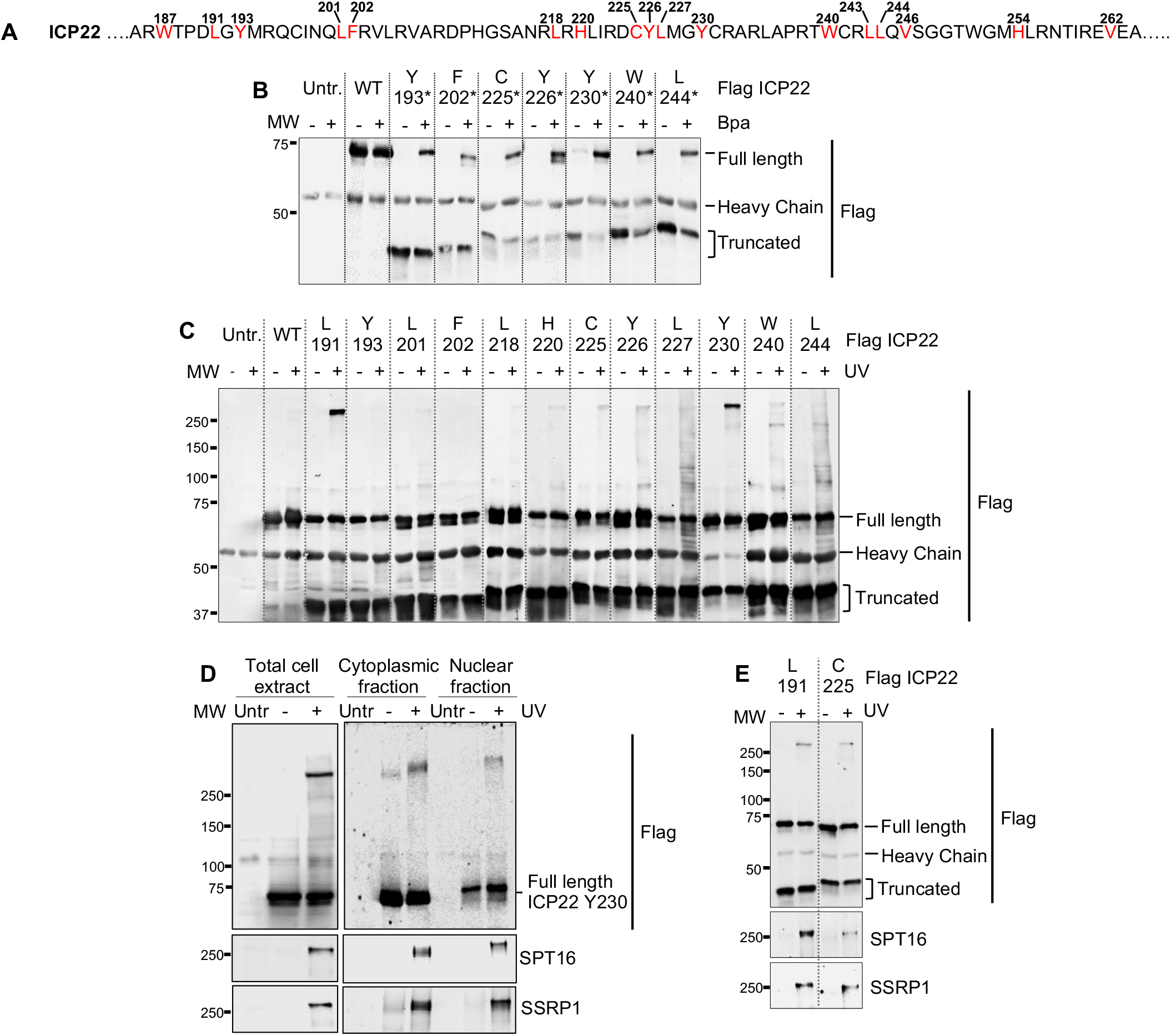
UV crosslinking reveals that ICP22 interacts directly with the FACT complex. A. Amino-acid sequence of ICP22 showing the positions at which a TAG stop codon is introduced and Bpa is incorporated within ICP22. B. HEK293 cells were cotransfected with wild-type ICP22 cDNA (WT) or ICP22 cDNA with a TAG stop codon replacing different amino acid codons and suppressor tRNA/Bpa synthetase pair in the absence or presence of photo-crosslinkable Bpa. Flag immunoprecipitation was performed from total cell extracts and separated by SDS-PAGE. Western blots were probed with an anti-Flag antibody. C. Flag immunoprecipitation of ICP22^Bpa^ from cell extracts after cells were UV irradiated alive (+) in chilled PBS or not (-). Western blot was probed with an anti-Flag antibody. D. HEK293 cells were cotransfected with ICP22 cDNA with a TAG stop codon replacing Y230 and suppressor tRNA/Bpa synthetase pair in the presence of photo-crosslinkable Bpa, and UV-irradiated alive (+) in ice-cold PBS. Flag immunoprecipitation was performed from total cell extracts, cytoplasmic and nuclear fractions and separated by SDS-PAGE. Western blots were probed with antibodies against Flag and FACT complex subunits SPT16 and SSRP1. E. HEK293 cells were cotransfected with ICP22 cDNA with a TAG stop codon replacing L191 or C225 and suppressor tRNA/Bpa synthetase pair in the presence of photo-crosslinkable Bpa, and UV-irradiated alive (+) in ice-cold PBS. Flag immunoprecipitation was performed from total cell extracts and separated by SDS-PAGE. Western blots were probed with antibodies against Flag and FACT complex subunits SPT16 and SSRP1.

To generate ICP22-cellular protein crosslinks, transfected cells were UV-irradiated. Benzonase^®^ nuclease-treated cell lysates were immunoprecipitated with anti-Flag beads and analysed by western blot using an anti-Flag antibody. A high molecular weight band (>250kDa) was strongly detected upon UV irradiation of ICP22^Bpa^ L191 and Y230 compared with other mutants and Flag-ICP22^WT^ (Fig. 3C). Several other crosslinks of molecular mass between 100-250 kDa were also observed from ICP22^Bpa^ L227, W240 and L244. We were particularly interested in ICP22^Bpa^ Y230 that gives efficient crosslinking to large cellular proteins. To identify the unknown large crosslink product(s), UV-irradiated samples were silver-stained, and mass spectrometry analysis was performed from a gel slice containing proteins with a molecular mass of approximately 250 kDa, with controls of untransfected cells or samples not subjected to UV-irradiation (Supplementary Fig. S3(A)). The highest score for the band above 250kDa corresponds to ICP22 having incorporated Bpa in peptide 224-DCYLMGYCR-232 that includes Y230 (Supplementary Table 3). Several other proteins present in the crosslinked sample were identified with at least 30-fold higher intensity than in control samples from cells not exposed to Bpa or not irradiated (Supplementary Fig. S3(B) and Supplementary Table 4). Proteins that are identified by at least two unique peptides belong to three classes: a) chaperones (HSP90 alpha and beta, HSC71, HSP70.1A, HSP70.4, HSP70.6, FAM10A5, stress–induced phosphoprotein and ubiquitin modifier); b) transcription factors (SPT16, SSRP1, TFIIIC.1) and transcript processing factors (PRPF8, PARP1, SFPQ, hnRNP M, THO complex subunit, THOC2).

Interestingly, both FACT subunits, SPT16 and SSRP1 are specifically detected after crosslinking. Coimmunoprecipitation of ICP22^Bpa^ Y230 upon UV irradiation was performed from total cell lysates, nuclear and cytoplasmic fractions using anti-Flag beads. Western blot using anti-Flag antibody and antibodies to both FACT subunits was able to detect the large crosslink in both cellular compartments, thus confirming the ICP22xFACT crosslink (Fig. 3D). Residues ICP22^Bpa^ L191 and C255 also crosslink to SPT16 and SSRP1 (Fig. 3E), indicating that several residues within the core conserved region of ICP22 are in contact with the FACT complex. The ICP22^Bpa^ Y230 crosslink was tested by western blot using antibodies against PARP1, PRPF8, SFPQ, TRIM28 and THOC2. However none of them was detected as part of the crosslinked product (data not shown).

### VP16 relieves ICP22-mediated inhibition

The structure of ICP22^CCR^ may provide insight into how ICP22 inhibits crucial host factors involved in host transcription elongation. However structural information is lacking at present. Sequence alignment and homology modelling suggest that the ICP22^CCR^ could adopt a helix-turn-helix conformation that closely resembles the concave surface of the VP16 core region (Fig. 4A). VP16 interacts with the conserved cyclin box domain of Cyclin T1 and increases P-TEFb recruitment to the viral promoter through recognition of the TAATGARAT motif of immediate early gene in association with HCF-1 and Oct1 (5, 6, 32). To investigate the role of the core region of VP16, we expressed Flag-tagged full-length VP16 and its core region (labelled CORE; residues 177-240) in HEK293. Coimmunoprecipitation followed by western blot indicates that the VP16^CORE^ interacts with P-TEFb (Fig. 4B). As VP16 competes with ICP22 for interaction with P-TEFb (6), we asked whether VP16^CORE^ is responsible for competition. Full length ICP22 and VP16 (Fig. 4C) or ICP22^CCR^ and VP16^CORE^ (Fig. 4D) were expressed individually or together and the level of Ser2P pol II CTD and CDK9 pThr186, which is a mark of CDK9 enzymatic activity, were analysed by western blot. As expected, full-length ICP22 induces loss of Ser2P pol II CTD, while VP16 does not. When both proteins are co-expressed, the level of Ser2P is restored, indicating that VP16 abrogates ICP22 inhibition of P-TEFb. Similarly, ICP22 causes a marked decrease in CDK9 pThr186 whereas VP16 does not and expression of VP16 with ICP22 restores CDK9 Thr186 phosphorylation.

**Fig. 4:**
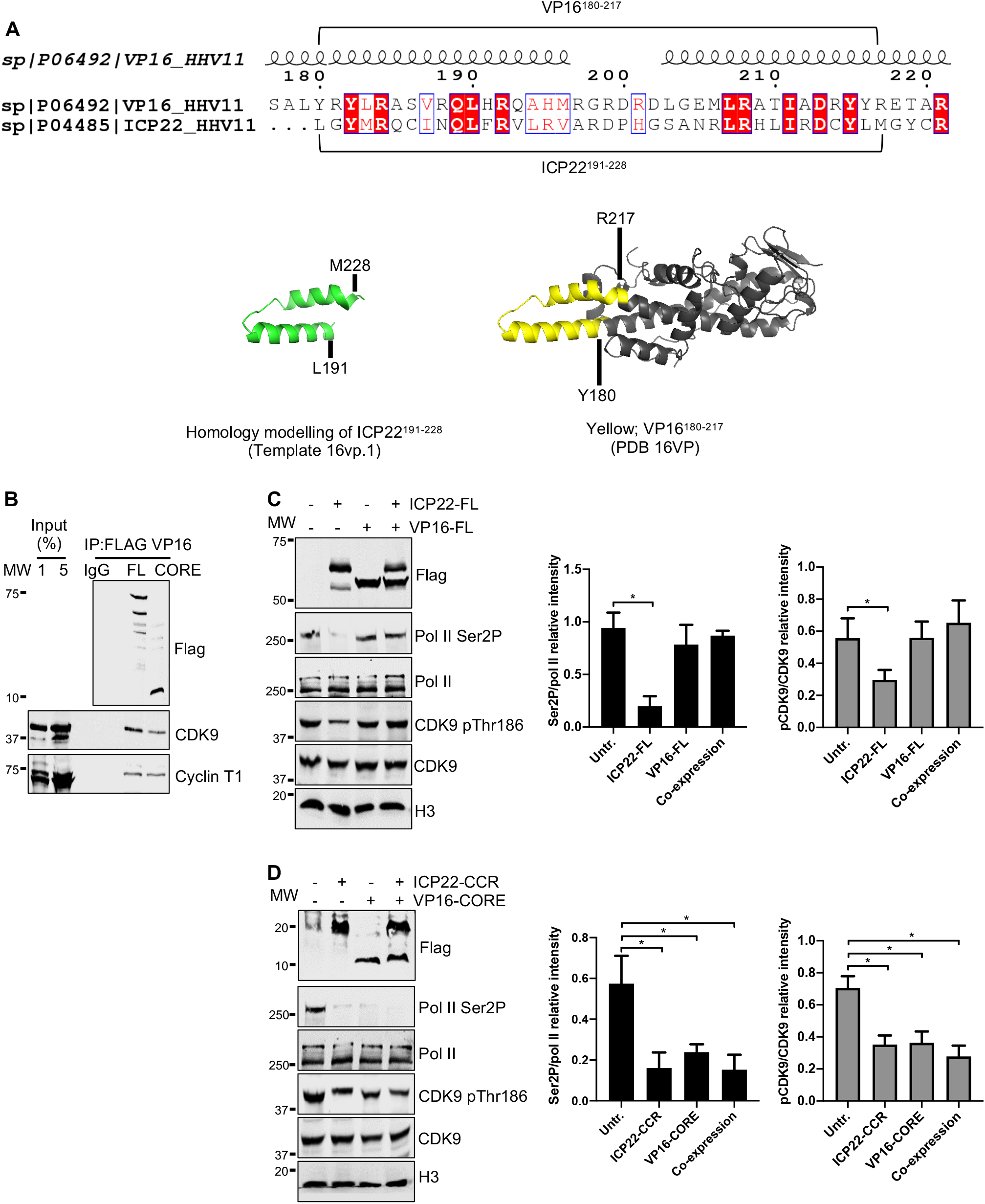
VP16 relieves ICP22-mediated inhibition. A. (Top) The pairwise alignment of VP16 and ICP22 was generated using ESPript. The secondary structure of VP16 is placed above the alignment. Identical amino acids are in red boxes and those that share similar properties of side chains are indicated by red letters. (Bottom) Homology modelling using ICP22 or VP16 as templates. B. HEK293 cells were transiently expressed with Flag-VP16, either full length (FL) or the CORE (residue 177-240). Cells were lysed, nuclear lysates were carefully separated for IP assays using control IgG or anti-Flag beads. Copurified proteins were determined by western blot using the antibodies noted at the right. C. HEK293 cells were transfected with plasmids expressing Flag-ICP22^FL^ or Flag-VP16^FL^ alone or together for 42 h. Nuclear lysates were analysed by western blot with Ser2P, pol II, CDK9 pThr186, CDK9 and H3 antibodies. H3 serves as a loading control. The relative intensity of protein signal of Ser2P/pol II and CDK9 pThr186/CDK9 were calculated from three independent experiments. Error bars indicate SEM and asterisks indicate the statistically significance of a change in signal compared with untransfected cells (Untr) (p<0.05). D. HEK293 cells were transfected with plasmids expressing Flag-ICP22^CCR^ or Flag-VP16^CORE^ alone or together for 42 h. Nuclear lysates were analysed by western blot with Ser2P, pol II, CDK9 pThr186, CDK9 and H3 antibodies. H3 serves as a loading control. The relative intensity of protein signal of Ser2P/pol II and CDK9 pThr186/CDK9 were calculated from three independent experiments. Error bars indicate SEM and asterisks indicate the statistically significance of a change in signal compared to untransfected cells (Untr) (p<0.05).

Interestingly, the VP16^CORE^ induces loss of Ser2P pol II CTD in the absence or presence of ICP22^CCR^, indicating that VP16^CORE^ inhibits P-TEFb. The inhibition is also reflected by reduction of CDK9 pThr186 when ICP22^CCR^ or VP16^CORE^ are expressed individually or together. This result suggests that VP16^CORE^ and ICP22^CCR^ may inhibit the activity of P-TEFb and possibly other CTD kinases in a similar way.

Thus, VP16 can overcome the effect of ICP22 on pol II CTD kinases but VP16^CORE^ appears to function like the ICP22^CCR^. Other regions of VP16 must therefore be involved in relieving the inhibition; for example, the VP16 transactivation domain (TAD).

Alternatively, host cell proteins that interact with VP16 may be needed to reverse inhibition of pol II CTD kinases. The core region of VP16 may perform a critical role by competing P-TEFb away from ICP22 to allow delivery to viral genes for activation.

## Discussion

The HSV-1 viral genes are arranged in the same way as host-cell pol II-transcribed genes, with TATA box-containing promoters, introns and polyadenylation signals (33-35). However, after infection of cells, HSV-1 needs to downregulate expression of host genes while upregulating expression of the viral genome. HSV-1 infection induces the production of a modified species of pol II with Ser5P in the CTD but lacking Ser2P, indicating that transcription elongation is compromised through inhibition of Ser2 pol II CTD kinases like P-TEFb (28, 36). Previous studies, supported by the results presented here, indicate that ICP22 plays a major role in P-TEFb inhibition and the consequent inhibition of host-cell gene expression, which will in turn repress the expression of antiviral proteins as part of an immune evasion strategy (37-40). However, as P-TEFb activity is required for productive elongation of transcription of HSV-1 genes (41), the virus must have a sophisticated strategy to inhibit elongation only of host-cell genes. In order to more fully understand how this is achieved, we have used several complementary approaches to study the interaction of ICP22 with host-cell proteins.

A time course of transient expression of ICP22 in HEK293 revealed that ICP22 can cause an identical pattern of inhibition of transcribing pol II to treatment of cells with small molecule CDK9 inhibitors, like DRB (27). However, small molecule CDK9 inhibitors and ICP22 may also inhibit other CTD kinases. Using an unbiased approach of mass spectrometry coupled with western blot of the cellular proteins copurified with ICP22^CCR^, we identified CDK9 and other transcriptional kinases, CDK12, CDK11B and CDK13 that interact with and may be inhibited by ICP22. CDK12 is of particular interest as it is another elongation-phase kinase, which has been reported as being sensitive to DRB (22, 42). Most transcriptional CDKs have high sequence similarity in the kinase domain and a similar structural arrangement, with ATP-binding sites between the N- and C-terminal lobes, a hinge region that connects the two lobes, a cyclin-binding domain, and a similar activation segment. Generally, the cyclin interacts with the N-terminal lobe of the CDK while the C-terminal lobe of the CDK contains the motif recognising its substrates (25, 43). Because CDK12/Cyclin K have a similar structural arrangement to CDK9/Cyclin T and phosphorylates Ser2 pol II CTD (23, 42), we proposed, based on our results, that ICP22 may also inhibit CDK12/Cyclin K and the inhibition may have functional consequences for host transcription elongation during HSV-1 infection. In agreement with previous studies (15, 30), we found no interaction between ICP22 with CDK7, a component of the general transcription factor TFIIH and a CTD kinase which mainly targets Ser5 of the pol II CTD. CDK7 has a critical role early in the transcription cycle and Ser5P predominates on promoter-proximal pol II (44).

The interaction of ICP22 and P-TEFb-associated factors, including the FACT complex, further validates the role of ICP22 in regulating pol II-mediated transcription elongation of host-cell genes. FACT is a histone chaperone that facilitates transcription by disruption of the DNA-histone interface which then enables the passage of elongating pol II through nucleosomes (20, 45). This function of the FACT complex is also involved in DNA replication and repair (46, 47). No histones are found associated with HSV DNA in the virion and the nucleosomes are deposited on HSV DNA during the earliest stage of infection allowing circularization of the originally linear viral genome (48). Thus, cellular histone chaperone proteins are important for the accumulation of nucleosomes on HSV DNA before and during the different phases of viral gene expression (49). It is therefore likely that the association of the FACT complex with ICP22, one of the immediate early proteins, plays a very important role early in infection. This notion is further supported by a previous study indicating that ICP22 recruits FACT for efficient transcription elongation of viral genes (14). As ICP22 is associated with host genes (11) and does not play an essential role in viral DNA replication (14, 50, 51), we postulate that the interaction of ICP22 with the FACT complex also take place on host genes to help repress FACT-dependent transcription.

We show that multiple residues within the core conserved region of ICP22 crosslink to FACT in living cells, which implies that the association is stable, and both FACT complex subunits were associated with ICP22 after purification under stringent conditions. Both the SPT16 and SSRP1 subunits of FACT share general structural features such as a dimerization domain nearby the N-terminal region, a middle domain and a short, highly acidic C-terminal region, that are responsible for flexible interaction with histones and nucleosomal DNA (Fig. 5A; domain structure) (45, 52, 53). The N-terminal region and middle domain of the FACT heterodimer offer an interaction interface separate from the C-terminus, which binds the nucleosome (Fig. 5A; FACT-nucleosome diagram). The predicted structure of ICP22^CCR^ may therefore be positioned close to the dimerization domain of SPT16 and SSRP1. It is also possible that ICP22 crosslinks to the C-terminus of histone-free FACT complex, thus impairing its essential role as a histone chaperone. The sequence alignment of Cyclin T1 and N-terminal regions of SPT16 and SSRP1 reveal that residues within these ICP22-interacting proteins are poorly conserved (Supplementary Fig. S4), suggesting the formation of unique ICP22/P-TEFb and ICP22/FACT complexes. Future studies including structural analysis will be necessary to test this hypothesis.

**Fig. 5:**
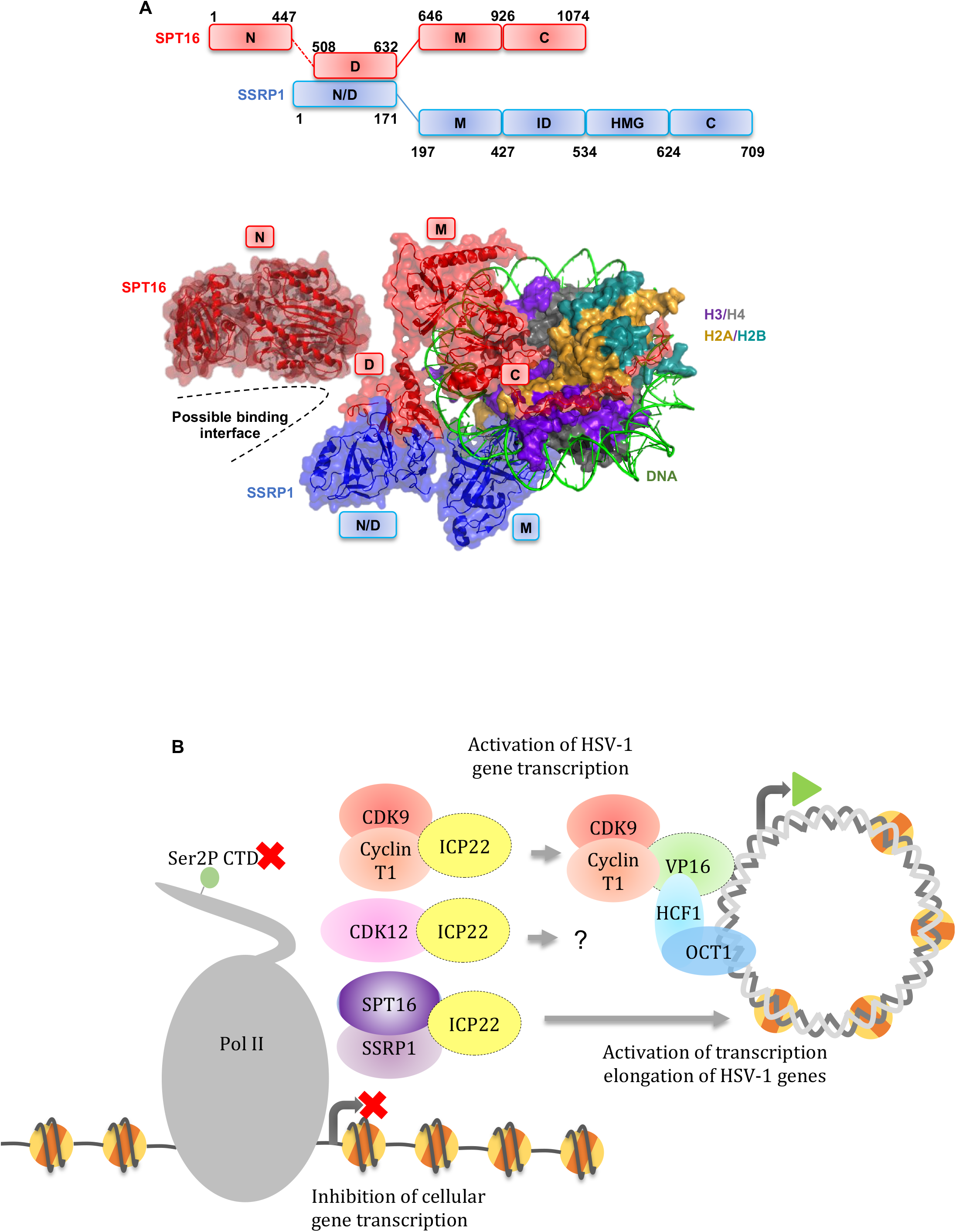
ICP22 interacts with key elongation regulators leading to repression of cellular genes. A. (Top) Domain organization of FACT heterodimer SPT16 and SSRP1. N: N-terminal domain; D: Dimerization domain; M: Middle domain; C: C-terminal domain; ID: intrinsically disordered domain; HMG; HMG box domain. (Bottom) Structure of FACT complex subunits and nucleosome (PDB 6UPL (52); N-terminal SPT16 PDB 5XM2 (59)) generated with PyMOL. A possible interaction interface is denoted by dashed lines. B. Current working model. HSV-1 ICP22 and VP16 cooperatively regulate transcription elongation complexes. ICP22 affects cellular transcription elongation by inhibiting Ser2 pol II CTD kinase P-TEFb and VP16 reverses repression of P-TEFb to enhance viral transcription. The function of other transcription elongation regulators including additional Ser2 pol II CTD kinases such as CDK12 and the histone chaperone FACT complex might also be affected on host cell genes.

Surprisingly, our amber suppression-mediated photocrosslinking experiments did not detect a stable association between ICP22 and P-TEFb. However, this approach does not necessarily reflect the inability of ICP22 to interact with host proteins like P-TEFb, and selection of other residues to modify or alternative photocrosslinkers such as arylazide- or diazirine-containing amino acids may be necessary to detect this interaction (54).

HSV-1 VP16 is widely studied as a potent transactivator in the early phase of infection. VP16 is a type IIB activator which strongly stimulates initiation as well as elongation of transcription and phenylalanines within its C-terminal activation domain are key for VP16 activity (32, 55). ICP22 has no apparent DNA binding domain and does not associate with VP16 (6). However, our results indicate that VP16 can reverse ICP22-dependent inhibition of CTD kinases, indicating that there is crosstalk between these proteins in the cell. As the VP16^CORE^ has potential structural homology to the ICP22^CCR^ and can inhibit phosphorylation of Ser2 pol II CTD, the VP16^CORE^ may be able to compete with ICP22 for P-TEFb to bring it to the viral genes. On the basis of these new and previous findings, we propose a working model where ICP22 represses the activity of the Ser2 pol II CTD kinases P-TEFb and/or CDK12 and interacts with the FACT complex, resulting in promoter-proximal stalling and premature termination of elongating pol II on cellular genes (Fig. 5B). VP16 is able to relieve the repressive action of ICP22 and redirects these cellular resources for transcription of the viral immediate early genes.

Collectively, our results suggest that ICP22 selectively inhibits transcription elongation, at least in part by association with Ser2 pol II CTD kinases P-TEFb and/or CDK12 and the FACT elongation complex. Exactly how ICP22 orchestrates distinct interactions with the individual components of transcription elongation complexes leading to downregulation of host gene transcription and activation of viral genes by VP16 at a particular stage of HSV-1 infection remain to be determined.

## Materials and methods

### Cell culture and transfection

HEK293 (ATCC^®^ CRL-1573™) were maintained in DMEM medium supplemented with 10% foetal calf serum, 100U/ml penicillin, 100 μg/ml streptomycin, 2 mM L-glutamine at 37°C and 5% CO_2_. Cells were transfected with mammalian expression plasmids using jetPRIME® transfection reagent (Polyplus) as per the manufacturer’s recommendations.

### Plasmid construction

The mammalian expression plasmid encoding Myc-ICP22^CCR^ was described previously (11). The mammalian expression plasmids encoding Flag-ICP22 (full length and CCR) and Flag-VP16 (full length and CORE) were generated using restriction cloning into pcDNA3.1. The pcDNA3.1-Flag-ICP22 plasmids carrying an amber stop codon at selected positions were generated by standard PCR-based site-directed mutagenesis. The bacterial expression plasmid for His-ICP22^CCR^ and His-IpgC (kindly provided by Susan Lea) were generated using restriction cloning into pET28a, and GST-CDK9^FL^ and GST-Cyclin T1^1-266^ were cloned into pGEX-6P-1.

### Chromatin immunoprecipitation (ChIP)

ChIP was performed as described previously (27), using antibodies against control IgG (sc-2027; Santa Cruz), Pol II N20 (sc-899-X; Santa Cruz) and Ser2P (ab5095; Abcam). ChIP samples were evaluated by real-time qPCR (qRT-PCR) using QuantiTect SYBR Green PCR master mix in a Rotor-Gene 3000 cycler (Corbett Research). The sequences of primers used for qRT-PCR are listed in Supplementary Table 5. Immunoprecipitated and control DNA samples were analysed in triplicate reactions. The ChIP signal is presented as a percentage of input DNA after removing the background signal from DNA immunoprecipitated with IgG. Data represent the mean ± SEM (standard error of the mean) of three independent experiments and p-values were calculated by two-tailed unpaired t-test in which p<0.05 was considered to be statistically significant.

### Coimmunoprecipitation (CoIP)

CoIP was performed from a 15 cm dish of HEK293 transfected with mammalian expression plasmids. Cells were harvested in ice-cold PBS supplemented with protease inhibitor cocktail. The cell pellet was lysed in lysis buffer (10 mM HEPES, 10 mM KCl, 150 mM NaCl, 0.1 mM EDTA, 0.1 mM EGTA, 0.5% NP-40 and protease inhibitor cocktail). The suspension was centrifuged at 13,000 g for 15 min at 4°C and the nuclear pellet was lysed in IP buffer (10 mM HEPES, 150 mM NaCl, 0.1 mM EDTA, 0.1 mM EGTA, 0.5% NP-40, 0.1% SDS, 0.1% sodium deoxycholate and protease inhibitor cocktail). The lysate was sonicated in chilled water bath and treated with 25 units of Benzonase (Merck Millipore) for 1 h at 4°C. After centrifuging for 15 min at 13,000 g at 4°C, the soluble supernatant was transferred into a fresh tube. Pre-equilibrated anti-Flag M2 beads (Sigma Aldrich) were added into the supernatant and incubated overnight at 4°C on a wheel. The beads were washed three times with IP buffer.

### Western blot

Proteins in Laemmli sample buffer were boiled for 10 min at 95°C and electrophoresed in 8% or 12% SDS-PAGE gels to resolve proteins ≥ 20kDa. Smaller proteins were resolved by SDS-PAGE with Tris Tricine-SDS running buffer (56). Proteins were electrotransferred to 0.2 μm or 0.45 μm nitrocellulose. Blots were blocked with 5% BSA prepared in Tris-buffered saline (20 mM Tris-HCI pH 7.6, 137 mM NaCl, and 0.2% Tween 20) for 1 h and incubated with primary antibodies overnight at 4°C. The following primary antibodies were used: Pol II (NBP2-32080; Novus Biologicals), Ser2P (5095; Abcam), CDK9 C20 (sc-484; Santa Cruz), CDK9 pThr-186 (25495; Cell Signalling), CDK12 (A301-679A; Bethyl Lab), CDK12 (LS-C288466; LSBio), CDK12 2F6 (WH0051755M2; Sigma), CDK13 CDC2L5 (A301-458A; Bethyl Lab), CDK11 (A300-310A; Bethyl Lab), CDK8 (ab85854; Abcam), CDK10 C19 (sc-51266; Santa Cruz), CDK7 C19 (sc-529; Santa Cruz), Cyclin T1 H245 (sc-10750; Santa Cruz), Cyclin K G11 (sc-376371; Santa Cruz), Cyclin L1 (PA540668; Life Technologies), PAF1 (NB600-274; Novus Biologicals), LEO1 (A300–175A; Bethyl Lab), CDC73 (A300–170A; Bethyl Lab), SPT16 (607002; Biolegend), SSRP1 (609702; Biolegend), NELFE H140 (sc32912; Santa Cruz), MEPCE (14917-1-AP; Proteintech), SPT5 D10 (sc-390961; Santa Cruz), Myc (ab9132; Abcam), histone H3 (ab1791; Abcam), Flag M2 (F1804; Sigma Aldrich), 6xHis (ab18184; Abcam) and GST (27-4577-01; GE Healthcare). Secondary antibodies were purchased from Li-cor Biosciences; IRDye® 800CW Goat anti-Mouse IgG (925-32210), IRDye® 680RD Goat anti-Mouse IgG (925-68070), IRDye® 800CW Goat anti-Rabbit IgG (925-32211), IRDye® 680RD Goat anti-Rabbit IgG (925-68071), IRDye® 800CW Donkey anti-Goat (925-32214), IRDye® 680RD Donkey anti-Goat IgG (925-68074). Blots were visualized on an Odyssey CLx Imaging System (Li-cor Biosciences) and quantification of the protein intensity was performed with Image Studio Lite software.

### Pulldown of bacterially-expressed recombinant proteins

Recombinant proteins were expressed in *Escherichia coli* RosettaTM (DE3) (Millipore) by induction overnight with 0.5 mM isopropyl β**-**D**-**1-thiogalactopyranoside (IPTG) at 18 °C. Lysis buffer (50 mM Tris-HCl pH 8, 150 mM NaCl, 1 mM EDTA, 1 mg/ml lysozyme, 0.2% Triton X-100 and protease inhibitor cocktail) was used to purify His-tagged recombinant proteins and lysis buffer (20 mM Tris-HCl pH 8, 150 mM NaCl, 1 mM EDTA, 0.5% NP-40, mM dithiothreitol, 0.2 mM phenylmethylsulfonyl fluoride (PMSF) and protease inhibitor cocktail) was used to purify GST-tagged recombinant proteins. The suspension was centrifuged at 14,000 g for 30 min at 4°C. His-tagged recombinant proteins were retained on nickel magnetic beads (VWR International) and GST-tagged recombinant proteins were retained on glutathione magnetic beads (Fisher Scientific). On-bead recombinant proteins were washed with the respective lysis buffer. GST-tagged recombinant proteins were eluted with 10 mM reduced glutathione for pulldown assay with His-tagged recombinant proteins. For pulldown assay with HeLa nuclear extracts (Ipracell), the extracts were diluted 1:1 with IP buffer and precleared by incubating with nickel magnetic beads at 4°C on a wheel for 2 h. On-bead His-tagged recombinant proteins were added to the extract and incubated overnight. Bound proteins were washed three times with IP buffer.

### Photocrosslinking in live mammalian cells

UV-crosslinking followed the previously-described protocol (57) with minor changes. HEK293 cells were co-transfected with plasmids encoding suppressor tRNA (tRNA^Sup^) and aminoacyl tRNA synthetase, and mutated derivatives of pcDNA3.1-Flag-ICP22 in medium supplemented with p-benzoylphenylalanine (Bpa) (IRIS Biotech) at a final concentration of 2 mM. After 48 h, the medium was replaced with 5 ml PBS. Cells which were still attached to the plates were irradiated at 365 nm UV (4 J/cm^2^) (UVP 95-0007-06 UV lamp) for 45 min on ice. Cell pellets were resuspended in lysis buffer (10 mM HEPES, 10 mM KCl, 200 mM NaCl, 0.1 mM EDTA, 0.1 mM EGTA, 0.5% NP-40 and protease inhibitor cocktail). The suspension was centrifuged at 13,000 g for 15 min at 4°C and the supernatant was collected. The nuclear pellet was lysed in stringent IP buffer (50 mM HEPES, 1 M NaCl, 1 mM EGTA, 1 M Urea, 1% sodium deoxycholate, 1% NP-40 and 0.1% SDS). After this point, the lysates were prepared for coIP using the coIP protocol detailed above. To prepare mass spectrometry-compatible silver stained gels, the crosslink samples separated by SDS-PAGE gel were fixed in 50% ethanol, 10% acetic acid and 0.01% formaldehyde for 1 h before washing with 5 μg/ml dithiothreitol for 1 h. After being rinsed in water, gels were stained in freshly prepared 0.1% silver nitrate for 1 h and developed in 3% sodium carbonate and 0.01% formaldehyde. The reaction was terminated by the addition of 50% ethanol and 10% acetic acid. Gel bands of the crosslinked products, and equal size bands from untransfected and non-irradiated cells were excised and analysed by mass spectrometry.

### Mass spectrometry

Mass spectrometry analysis was performed by the proteomic facilities at the University of Oxford or Queen Mary University of London, essentially as described (58). Protein samples were treated with in-gel or on-bead trypsin digestion, and the purified proteolytic peptides were analysed by nano liquid chromatography with tandem mass spectrometry (LC-MS/MS) using a Q Exactive Plus Hybrid Quadrupole-Orbitrap Mass Spectrometer (Thermo Fisher Scientific). Data were searched using Mascot-driven database searches and label-free quantitative proteomics data were analysed based on the features from fragment-ion analysis (MS2). Protein and peptide identifications were filtered at 1% peptide-level false discovery rates and required at least two unique peptides per protein.

## Availability of data and material

The authors declare that all relevant data supporting the findings of this study are available within the article and its supplementary information files.

## Code availability

Not applicable

## Author’s contribution

S.M supervised the research and interpreted data. O.B provided materials, supervised UV crosslinking experiments and helped with data interpretation. N.C.A carried out analysis of protein structures. N.F.I produced materials, performed all of the experimental work and analysed data. N.F.I and S.M wrote the paper with contributions from the other authors. We are grateful to Michael Tellier for critical reading of the manuscript.

---

**Supplementary Fig. S1.**
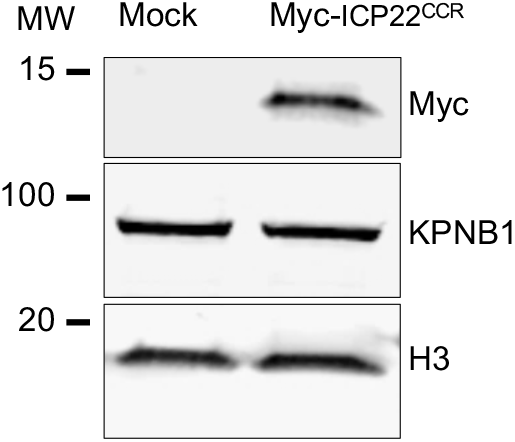
Western blot analysis of KPNB1 (Primary antibody A300-482A; Bethyl lab) in HEK293 cell extract after transfection of a vector expressing Myc-tag ICP22^CCR^. H3 is used as loading control.

**Supplementary Fig. S2.**
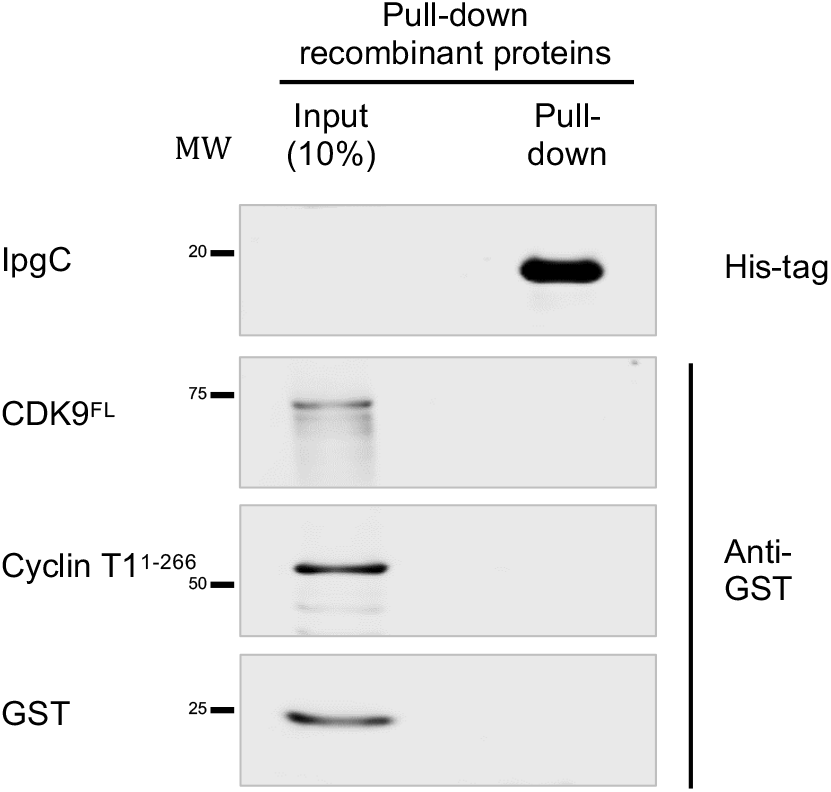
His-IpgC were bacterially expressed and immobilized on nickel magnetic beads. GST-Cyclin T1^1-266^ and GST-CDK9^FL^ were expressed separately in *E*.*coli*. Immobilized GST-Cyclin T1^1-266^ and GST-CDK9^FL^ were eluted with reduced glutathione, and incubated with on-bead His-IpgC. Both input and proteins retained on the beads after washes were subjected to western blot with GST and His-tag antibodies. Input represents 10% of that used for pull-down.

**Supplementary Fig. S3.**
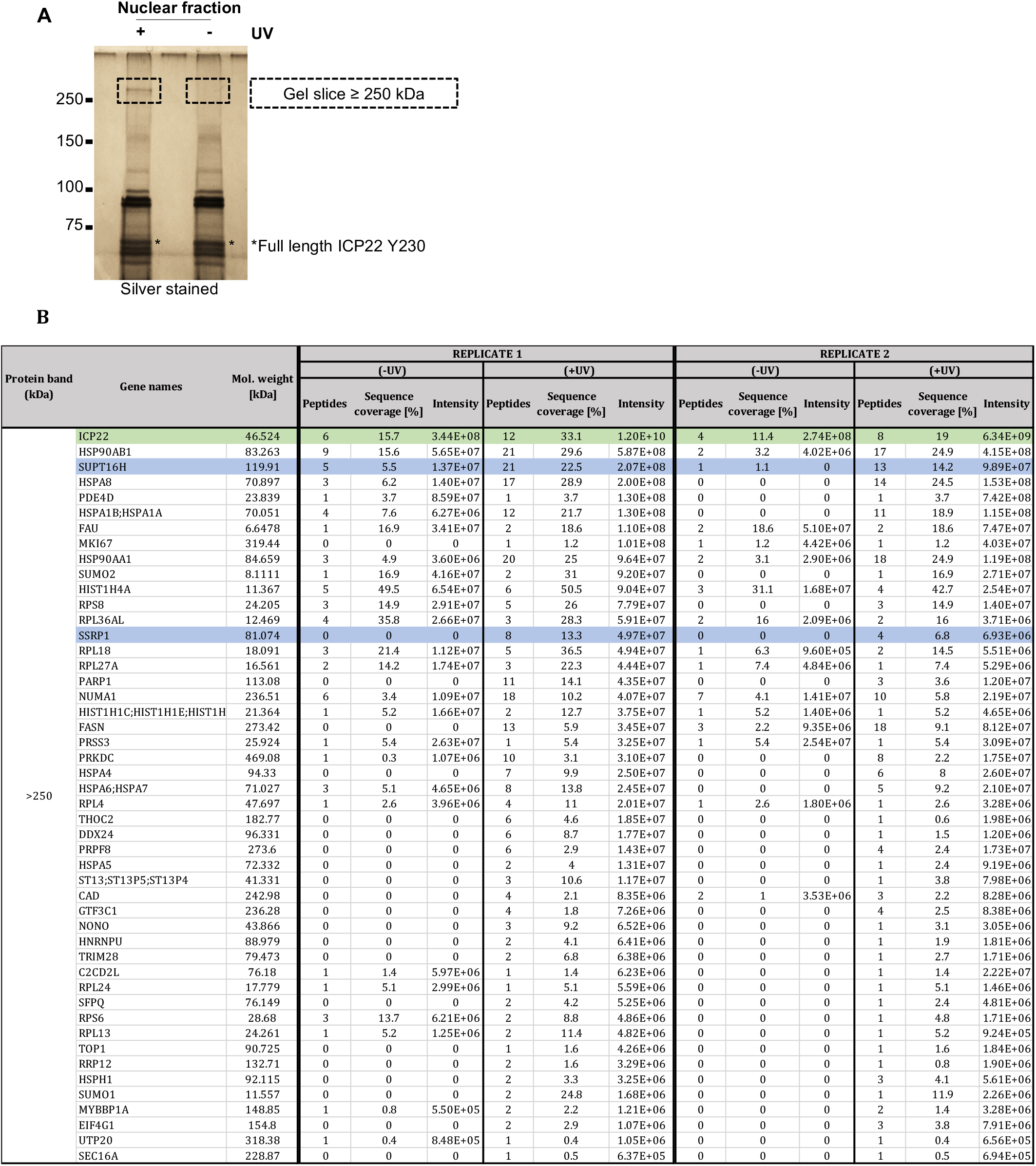
(A) Indicated protein bands (>250kDa) of silver-stained SDS-PAGE gel were excised, in-gel digested and analysed by mass spectrometry. (B) Mass spectrometric analysis of ICP22 (green shade), SPT16 (gene name; SUPT16H) and SSRP1 (blue shades) identified from the silver-stained gel slice of ≥250 kDa of no UV and UV-treated samples. Details of the proteomic data including the untransfected samples can be found in Supplementary Table 4.

**Supplementary Fig. S4.**
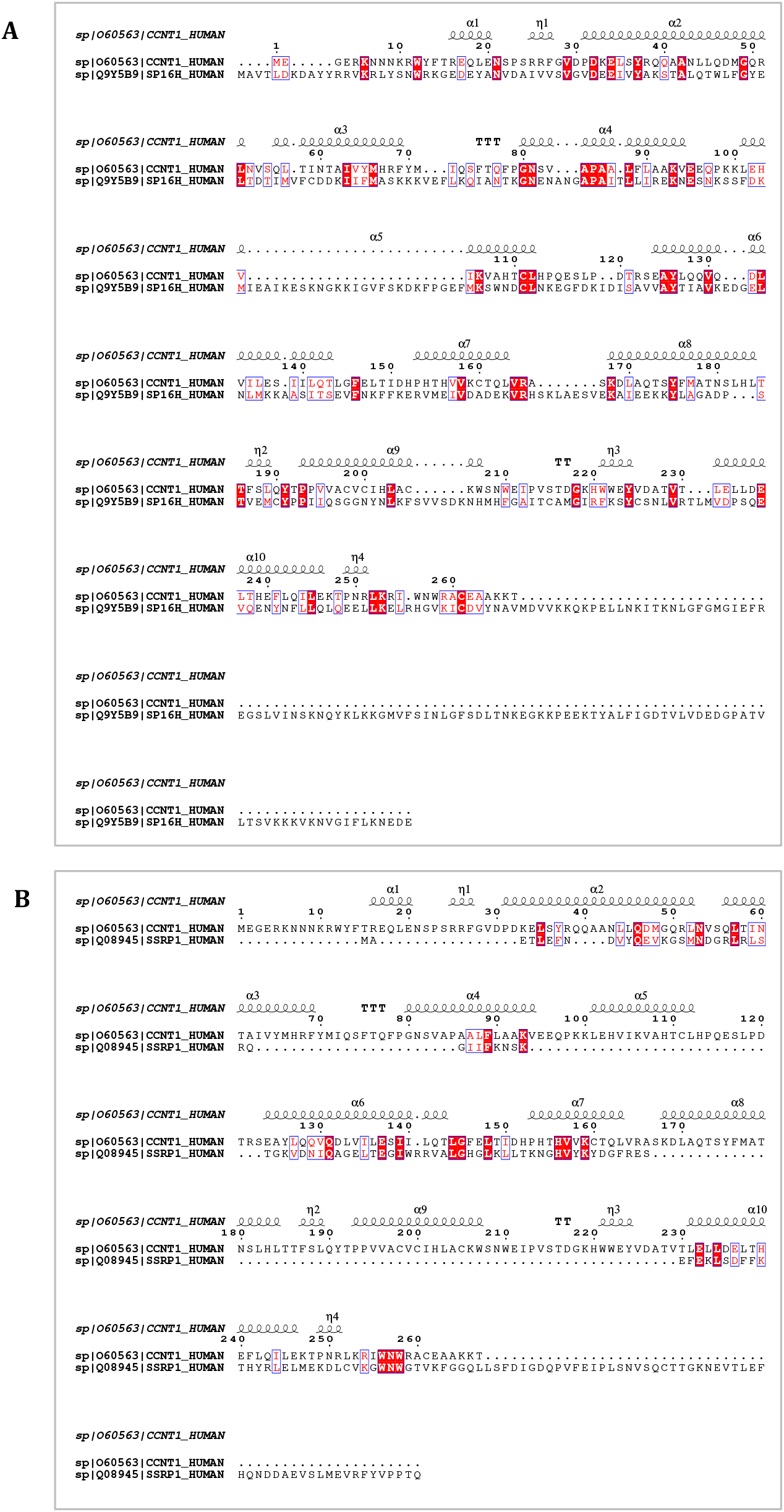
Sequence alignment of Cyclin T1 residues 1-266 (Uniprot ID O60563) and N-terminal region of (A) SPT16 (Uniprot ID Q9Y5B9) or (B) SSRP1 (Uniprot ID Q08945) showing minimal sequence similarity between the two known ICP22-binding complexes.

